# Proximity labeling reveals an extensive steady-state stress granule interactome and insights to neurodegeneration

**DOI:** 10.1101/152520

**Authors:** Sebastian Markmiller, Sahar Soltanieh, Kari Server, Raymond Mak, Wenhao Jin, Enching Luo, Florian Krach, Mark W. Kankel, Anindya Sen, Eric J. Bennett, Eric Lécuyer, Gene W. Yeo

## Abstract

Stress granules (SGs) are transient ribonucleoprotein (RNP) aggregates that form in response to proteotoxic stress. Although SGs are distinct from aggregates observed in neurodegenerative disorders, they share protein components. We used APEX-mediated proximity labeling combined with quantitative mass spectrometry and high-throughput imaging to identify >100 previously unknown SG proteins in human cells, about 10% of which localize to SGs in a cell type- or stress type-dependent manner. Supporting a link between SG proteins and neurodegeneration, we demonstrate aberrant SG composition and subcellular distribution in iPSC-derived motor neurons from ALS patients, and identify several known and previously unidentified SG proteins that modify toxicity of mutant FUS and TDP-43 overexpression in *Drosophila*. We show that even in an unstressed steady-state, SG proteins form a densely-connected protein interaction network (PIN) and propose a model in which existing RNPs coalesce rapidly into microscopically visible granules that can act as gateways to pathological protein aggregation.

**Highlights:**

- APEX proximity labeling of dynamic RNP granules identifies over 100 novel SG proteins
- SG proteins form a densely-connected protein interaction network in unstressed cells
- Systematic immunofluorescence analysis reveals stress- and cell type-specific SG composition
- ALS motor neurons contain SGs with distinct content and subcellular distribution

## Introduction

Cellular RNA molecules interact with a diverse array of nearly two thousand RNA binding proteins (RBPs) (Beckmann et al., 2015; Brannan et al., 2016; Gerstberger et al., 2014; Sheth and Parker, 2003) to form ribonucleoprotein particles (RNPs). The molecular makeup of RNPs is hypothesized to vary dynamically in response to changing cellular conditions as well as during different RNA processing requirements (Muller-McNicoll and Neugebauer, 2013). RNP assemblies span a range of sizes and compositional complexity. Nascent RNAs assemble into initial RNPs that can be tens of nanometers in size and can mature into larger RNP structures up to several microns in size. These large macromolecular structures are also known as RNP granules and represent microscopically visible and highly dynamic structures comprised of RNAs and proteins that function in diverse cellular pathways. Cajal bodies (CBs) which assist in the assembly of spliceosomal small nuclear RNPs (Gall et al., 1999) and processing bodies (PBs) that regulate RNA degradation and turnover in the cytoplasm are two examples of large nuclear and cytoplasmic RNP bodies, respectively (Ingelfinger et al., 2002; Sheth and Parker, 2003; van Dijk et al., 2002). In addition, there are cell type-specific RNP granules such as RNA transport granules in the axons and dendrites of neurons (Ainger et al., 1993; Knowles et al., 1996). While the above are all examples of RNPs that are present in healthy cells under ideal growth conditions, there is also a class of RNPs that is thought to form only in response to specific environmental insults. Exposure of cells to exogenous stresses can induce the rapid formation of large cytoplasmic RNP granules known as stress granules (SGs) (Kedersha et al., 1999; Protter and Parker, 2016). SGs are a particularly interesting type of RNP as their formation occurs concurrently with widespread RNA metabolism alterations, including global translation repression. Indeed, a primary function of SGs is thought to be sequestration and triage of untranslated mRNAs within stalled translation initiation complexes (Kedersha and Anderson, 2002).

Recently, there has been rapid progress in our understanding of SG biology. SGs appear to contain a dense stable ‘core’ whose formation requires an early ATP-dependent assembly step (Jain et al., 2016; Wheeler et al., 2016). Further SG growth is driven the accumulation of proteins containing intrinsically disordered regions (IDRs) and low complexity domains (LCDs). Once these proteins accumulate at high enough local concentrations, their IDRs and LCDs can undergo liquid-liquid phase transitions (LLPS), which then serve as the driving force for further reversible SG aggregation (Jain et al., 2016; Wheeler et al., 2016). In addition, several types of post-translational modifications (PTMs), including modification by O-GlcNAc (Ohn et al., 2008) and poly(ADP-ribose) (Leung et al., 2011) have been implicated in SG formation, but their role remains poorly understood.

In the past few years, SGs have been increasingly associated with human disease, most notably a subset of neurodegenerative disorders that are characterized by the presence of toxic insoluble protein aggregates. Amyotrophic Lateral Sclerosis (ALS) provides the most compelling link between SGs and neurodegeneration, as many of the proteins that compose pathological inclusions in patient cells are also found in transient SGs induced in normal cells. Moreover, ALS-linked mutations frequently occur within IDRs or LCDs of these proteins and these mutations are predicted to result in enhanced protein aggregation. These observations link the aberrant formation or persistence of SGs to the pathological insoluble protein aggregates characteristic to ALS.

Much progress has been made in understanding the molecular mechanisms of assembly and dispersion of SGs, but only few systematic approaches have been employed to comprehensively catalogue the protein and RNA content of SGs and other RNP granules (Buchan et al., 2013; Jain et al., 2016; Ohn et al., 2008). A recent proteomics analysis of isolated SG cores expanded the mammalian SG proteome and provided insights into molecular mechanisms of formation and resolution of these structures (Jain et al., 2016). However, it would be highly desirable to complement these efforts with *in vivo* approaches that address the question of potential artefactual loss or gain of SG-protein interactions following cell lysis.

In this study, we use a combination of APEX-mediated *in vivo* proximity labeling (Hung et al., 2014; Rhee et al., 2013) with quantitative mass spectrometry (MS) and an RBP-focused immunofluorescence approach to significantly expand the repertoire of known SG-associated proteins across several different cell types and stress conditions. We show that SG proteins form a dense protein interaction network in unstressed cells that is poised to enable rapid SG assembly in response to stress. In addition, we show that SG frequency and subcellular distribution are aberrant in motor neurons derived from stem cell models harboring ALS-associated mutations in a SG protein, hnRNP A2/B1. In addition, we show that several known and previously unknown SG components modify toxicity in Drosophila models of FUS and TDP-43-mediated degeneration. By systematically integrating our refined SG proteome with different neurodegeneration-relevant proteomics datasets we provide a rich resource for further investigations into the molecular mechanisms underlying SG formation and to elucidate the links between SGs and human disease.

## Results

### Cells expressing endogenously tagged G3BP1-APEX2-GFP allow for specific biotin labeling of stress granule proteins

To investigate the protein composition of SGs in living cells, we performed proximity labeling using an engineered ascorbate peroxidase (APEX2) fused to the well-characterized SG-associated protein G3BP1 (Figure 1A). We used CRISPR/Cas9-directed genome engineering to insert APEX2-GFP into the endogenous *G3BP1* locus in HEK293T cells (**Figure S1A**). The resulting G3BP1-APEX2-GFP fusion protein allows simultaneous visualization of SGs upon sodium arsenite (NaAsO_2_) exposure, as well as robust and rapid biotin labeling of SG-associated proteins only in the presence of biotin-phenol (BP) and upon a 60 second exposure to hydrogen peroxide (H_2_O_2_) (Figure 1B,C). We also established HEK293T cells with constitutive expression of a cytoplasmic-localized APEX2 (NES-APEX2-GFP) to serve as a specificity control (**Figure S1B**). Compared to G3BP1-APEX2-GFP expressing cells, NES-APEX2-GFP shows a diffuse cytoplasmic GFP signal that is unaffected by NaAsO_2_ treatment (Figure 1B). As expected, the overall pattern of protein biotinylation in NES-APEX2-GFP cells is unaffected by stress (Figure 1C), and remarkably similar to that of both unstressed and NaAsO2-treated G3BP1-APEX2-GFP cells. The majority of visible bands likely represent bystander biotinylation of highly abundant cellular proteins, underscoring the importance of a ratiometric quantitative proteomics approach. However, the observation that the biotinylation pattern of G3BP1-APEX2-GFP cells shows only subtle changes upon NaAsO_2_ treatment also suggests that despite the drastic re-localization of G3BP1-APEX2-GFP into SGs, the G3BP1 interactome remains relatively stable.

**Figure 1.**
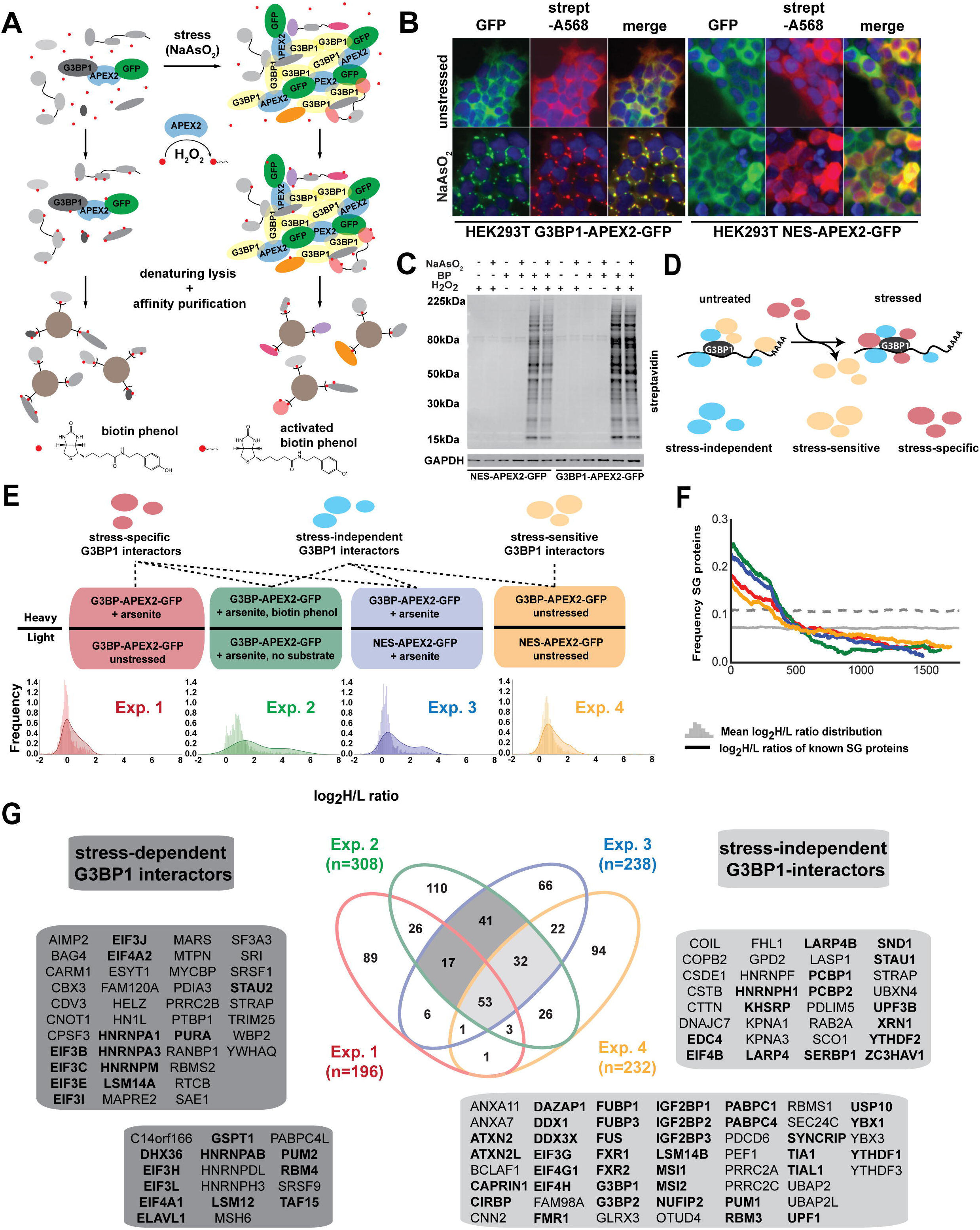
G3BP-APEX mediates specific biotinylation of stress granule-associated proteins. **A)** Schematic representing the use of proximity labeling by APEX2 to specifically tag stress-granule associated proteins with biotin. **B)** Streptavidin staining of unstressed (top panel) and sodium arsenite-treated (bottom panel) HEK293T G3BP-APEX2-GFP cells (left) and hPGK-NES-APEX2-GFP cells. **C)** Streptavidin-HRP western blot analysis of induced protein biotinylation in lysates from NES-APEX2-GFP and G3BP1-APEX2-GFP cells. **D)** Schematic depicting G3BP1 interactome changes upon stress treatment. **E)** Overview of four different experimental designs for detecting G3BP1-interacting proteins. Shown below the design are log2 heavy/light (H/L) ratio distributions of all proteins detected, overlaid with log2 H/L ratio distributions of known stress SG proteins detected in this experiment. **F)** Enrichment frequency distribution of known SG proteins in log_2_ H/L-ranked proteomics datasets. The frequency of known SG proteins was calculated on a rolling window basis with a window size of 500. The dashed line represents 1.5 times the background frequency of SG proteins across all detected proteins. **G)** Venn diagram showing overlap between hits from the four different experimental approaches and proteins found in the highlighted overlapping sets. Previously known SG proteins are highlighted in bold. See also Figure S1.

### Quantitative proteomics reveals increased biotinylation of known SG-associated proteins upon stress treatment

Since G3BP1 has an essential role in SG biology and robustly localizes to SGs, we reasoned that defining the interactome proximal to G3BP1 under stress conditions would approximate the SG proteome. However, little is known about the extent to which G3BP1 interactions change in response to cellular stress. To address this question, we next employed a SILAC-based quantitative proteomics approach with a series of experimental designs to systematically identify G3BP1-interacting proteins in both unstressed and stressed cells (Figures 1E and **S1C**). We hypothesized that there would be three classes of G3BP1-interacting proteins (Figure 1D): ***i)* stress-independent** interactors, such as USP10 and CAPRIN1 (Kedersha et al., 2016; Sowa et al., 2009), which associate with G3BP1 independently of stress; ***ii)* stress-dependent** partners that associate with G3BP1 only upon coalescence of large, microscopically-visible SGs that are induced by stress; and ***iii)* stress-sensitive** interactors whose association with G3BP1 is lost or weakened upon SG formation (Figure 1D).

To distinguish these different types of interaction partners, we pursued the four experimental schemes outlined in Figure 1E (upper panels). First, to identify stress-specific G3BP1 interactors, we characterized the biotinylated products obtained from stressed versus unstressed G3BP1-APEX2-GFP cells incubated with biotin-phenol (BP) (**Exp. 1**), which should enrich for proteins whose association with G3BP1 significantly increases or occurs only under stress. Next, to control for biotinylation reaction specificity, we contrasted lysates from stressed G3BP1-APEX2-GFP cells incubated with BP to lysates of an identical specimen for which the BP substrate was omitted (**Exp. 2**). Thirdly, to control for false-positive identifications resulting from diffuse cytoplasmic labeling by G3BP1-APEX2-GFP, we also compared lysates from stressed G3BP1-APEX2-GFP and NES-APEX2-GFP cells (**Exp. 3**). Lastly, to identify stress-independent G3BP1 interactors as well as stress-sensitive interactors, whose association with G3BP1 decreases upon stress, we profiled lysates from unstressed G3BP1-APEX2-GFP and NES-APEX2-GFP cells (**Exp. 4**).

For each experimental configuration, we conducted three replicate affinity-purifications of biotinylated proteins from mixed lysates using bead-conjugated streptavidin. Affinity-purified samples and the corresponding mixed input samples were analyzed by quantitative mass spectrometry. In total, we detected 1,006 proteins across all input samples and 1,774 proteins across all streptavidin enrichments (**Figure S1D**, **Table S1**), accounting for ~60% (136/236) of a manually curated list of 236 annotated SG proteins (**Table S2**).

To evaluate the power of each experimental design to identify known and novel SG proteins, we compared the enrichment of known SG proteins to the background distribution of all detected proteins (Figure 1E, lower panels). Known SG-associated proteins were clearly enriched in all four approaches, with the greatest shift in log2 H/L ratios detected when NaAsO_2_-treated G3BP1-APEX2-GFP lysates were compared to either the no-substrate control lysates or the NaAsO_2_-treated NES-APEX2-GFP lysates. Strikingly, enrichment of known SG proteins was far less pronounced when comparing stressed versus unstressed G3BP1-APEX2-GFP lysates. Moreover, we observed that even in the absence of cellular stress, known SG proteins are already significantly enriched in G3BP1-APEX2-GFP over NES-APEX2-GFP lysates. We conclude that the G3BP1 interactome does not change drastically upon exposure to stress, but rather, G3BP1 is already in close proximity to many SG-associated proteins in an unstressed steady state.

### G3BP1-APEX2-mediated biotinylation identifies SG proteins with high specificity

To identify proteins enriched for each experimental condition, we used the list of previously known SG-associated proteins (**Table S2**) as a benchmark for discovery and calculated their frequency distribution across the log2 H/L ratio-ranked lists of affinity-purified biotinylated proteins for each experiment (Figure 1F). For each ranked list, a cut-off was determined to be the point at which the frequency of known SG proteins in a moving window was 1.5 lower than the background frequency of known SGs. Proteins with log2 H/L ratios above the cut-off in at least two out of three replicates were conservatively considered significantly enriched (**Table S3**). In total, we identified 143 SG-associated proteins whose distribution across the different experimental designs are shown in Figure 1G. Of these, we designated the set of 70 proteins that were identified as significant hits in all experimental approaches where lysates from NaAsO_2_-treated G3BP1-APEX2-GFP cells were compared to other lysates as our highest confidence set of SG-associated proteins (Figure 1G). Indeed, 87% (61/70) of proteins in this list have either previously been shown to associate with SGs (n=47, highlighted in bold in Figure 1G) or have been indirectly linked to SGs, either through closely related family members or paralogs (HNRNPAB, HNRNPDL, HNRNPH3, PABPC4L, YBX3, YTHDF3), functional screening (RBMS1/2) (Couthouis et al., 2011), stress-associated post-translational modifications (SEC24C, MSH6) (Leung et al., 2011; Pleschke et al., 2000; Zachara et al., 2011) or through documented interactions with known SG proteins. For example, the DEAD-box helicase DDX1 is well known to localize to SGs and was recently shown to form an RNA transport complex with C14ORF166, FAM98A/B and RCTB (Akter et al., 2017; Perez-Gonzalez et al., 2014), all of which we have identified as previously unknown SG-associated proteins in our APEX approach. In addition, both UBAP2 and UBAP2L robustly interact with several key SG proteins including FUS, G3BP2, ELAVL1 and others (Havugimana et al., 2012; Sowa et al., 2009; Wang et al., 2015).

Interestingly, our high-confidence SG protein set also contains ANXA7, its closest paralog ANXA11 as well as PDCD6 and PEF1, which have been shown to interact with ANXA11 either as a PDCD6 homodimer or a PDCD6/PEF1 heterodimer (Rolland et al., 2014; Satoh et al., 2002). While none of these proteins had previously been implicated in SG biology, ANXA11 was recently shown to harbor ALS-associated mutations, some of which lead to abnormal protein aggregation (Smith et al., 2017). In conclusion, our APEX2-based proximity labeling approach successfully identifies known and previously undiscovered SG-associated proteins with relevance to neurodegenerative disease.

### G3BP1-APEX2-mediated biotinylation yields insights into the dynamic changes in protein interactions induced by stress

To address the question whether SGs represent a completely *de novo* transient assembly of previously unassociated proteins or whether cellular stress simply reinforces existing cellular interactions to a point where microscopically visible aggregates are observed, we examined the proportion of SG proteins that already interact with G3BP1 in the absence of stress. We found that about 40% (58/143) of SG-APEX hits either newly associated with G3BP1 and SGs or showed a significant increase in enrichment upon exposure to stress (Figure 1E). Consistent with the concept that stalled pre-initiation complexes accumulate in SGs and are thus in greater proximity to G3BP1 as a result of NaAsO_2_-induced eIF2α phosphorylation, we identified 9 (out of 12 detected overall) individual subunits of the EIF3 and EIF4 translation initiation factors as stress-dependent G3BP1 interactors.

Remarkably, however, we found that almost 60% (85/143) of APEX-identified SG proteins, including many of the most recognizable SG proteins (e.g. CAPRIN1, FMR1, PABPC1/4, TIA1, USP10), already interact with G3BP1 in the absence of stress (Figure 1E). This further supports our conclusion that the rapid emergence of microscopically visible SGs upon exposure to cellular stress is based on an existing network of cellular interactions. Interestingly, we also detect several proteins with functions in RNA decay (e.g. CSDE1, EDC4, HNRNPF, KHSRP, UPF3B, XRN1, YTHDF2, ZC3HAV1) among the stress-independent G3BP1-interactors, suggesting that mRNA decay factors also form part of a pre-existing protein-protein interaction (PPI) network that allows cells to rapidly adapt mRNA metabolism upon cellular stress.

### G3BP-APEX proximity labeling in human neural neuronal cells reveals cell-type and stress-specific SG proteins

We next wanted to test whether this approach could successfully be applied to more complex cell types with an emphasis on neuronal lineages, as they are the most relevant to neurological diseases characterized by aberrant SGs. We therefore used the CRISPR approach described above to generate a G3BP1-APEX2-GFP expressing human induced pluripotent stem cell (iPSC) line and differentiated it to neural progenitor cells (NPCs) (Reinhardt et al., 2013) (Figure 2A). G3BP1-APEX2-GFP robustly localizes to SGs upon NaAsO_2_ treatment in NPCs, and streptavidin staining overlaps well with the GFP signal (**Figure S2A**). We then performed quantitative proteomics experiments, comparing NaAsO_2_-treated and unstressed G3BP1-APEX2-GFP expressing NPCs, as well as NaAsO_2_-treated G3BP1-APEX2-GFP cells with and without substrate (Figure 2B). To compare the effects of different stressors, we carried out parallel experiments using thapsigargin, which robustly induces SGs via activation of the unfolded protein response. We identified 3,292 unique total proteins in streptavidin IPs from NPCs (**Figure S2B**) and analysis of log_2_ H/L ratio distributions as well as the enrichment of known SG proteins in the log_2_ H/L ratio-ranked mass spectrometry data (Figure 2C), gave similar results to those observed in HEK293T cells. Based on this, we employed a similar approach to identify SG-associated proteins and determined a log2 H/L ratio cutoff by calculating the point at which the frequency of known SG proteins in a moving window across a log_2_ H/L-ranked list of identified proteins was 2 times lower than the background frequency of known SGs (Figure 2C).

**Figure 2.**
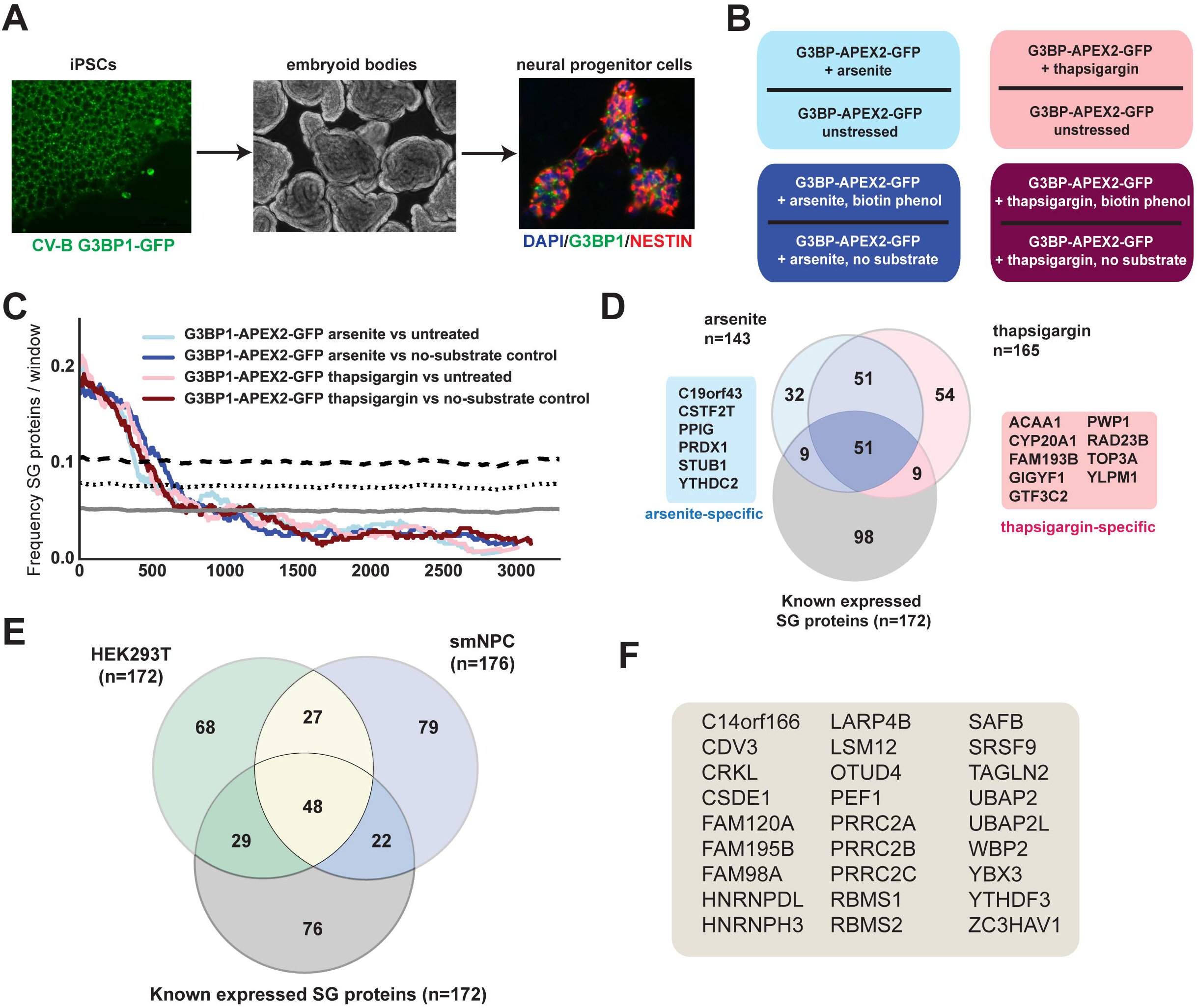
G3BP-APEX in HEK293T cells and NPCs identifies distinct but overlapping sets of SG-associated proteins. **A)** Overview of small molecule neuronal progenitor cell (smNPC) generation from induced pluripotent stem cells (iPSCs). **B)** Overview of experimental designs used in smNPCs. **C)** Enrichment frequency distribution of known SG proteins in log2 H/L-ranked proteomics datasets. The frequency of known SG proteins was calculated on a rolling window basis with a window size of 500. The dashed lines represent 1.5 and 2 times the background frequency of SG proteins across all detected proteins. **D)** Venn diagram showing the overlap between known SG proteins and APEX hits identified in cells stressed with NaAsO_2_ compared to thapsigargin. **E)** Venn diagram showing the overlap between known SG proteins and APEX hits identified in HEK293T cells and NPCs. **F)** Potential novel candidate SG proteins identified by G3BP-APEX in both HEK293T cells and NPCs. See also Figure S2.

Proteins with log_2_ H/L ratio satisfying this threshold in at least 3 out of 4 experimental conditions were considered to be SG-associated in NPCs. Using these criteria we identified 143 SG-associated proteins in NaAsO_2_-treated cells and 165 proteins in thapsigargin-treated cells (Figure 2D). Notably, 96% (137/143) of NaAsO_2_-induced and 95% (156/165) of thapsigargin-induced SG proteins were also found to be enriched in at least one experiment using the other stressor. While this suggests that SG composition is highly similar between these different stress conditions, we did identify 6 NaAsO_2_- induced and 9 thapsigargin-induced SG proteins that were stress-specific (Figure 2D). Altogether, we designated a total of 176 proteins as SG-associated in NPCs (**Table S4**), 40% (70/176) of which are known SG proteins, Lastly, we integrated the datasets from HEK293 cells and NPCs evaluate the extent to which SG composition is determined by cell type. For this, we used a set of 172 proteins in HEK293T cells that slightly expands on the 143 proteins described above by including an additional 29 proteins that were detected above threshold in at least 4 out of the 6 combined replicates from Exp.2 and Exp.3. (Figure 1E). We identified a set of 75 highest-confidence cell type-independent SG proteins (Figure 2E) 64% (48/75) of which are previously known SG proteins and 36% (27/75) are previously undiscovered SG proteins (Figure 2E, F).

### Post-translational protein modifiers localize to SGs but ubiquitylation is not acutely required for SG formation

We noticed that among the known and novel SG proteins identified across either of the two cell types were several proteins with roles in various post-translational protein modifications, including ubiquitylation (OTUD4, SQSTM/p62, STUB1, TRIM25, UBAP2L), sumoylation (SAE1) and other ubiquitin-like protein modifiers (GABARAPL2). We validated the proteomics data for several of these proteins by showing that they indeed co-localize with G3BP1-positive SGs (Figure 3A). While SGs have been shown to stain positive for ubiquitin (Kwon et al., 2007) and extensive poly-ubiquitylation is a hallmark of pathological protein inclusions in ALS (Arai et al., 2006; Neumann et al., 2006), it is still unknown whether active ubiquitylation is in fact required for SG formation. To test this, we treated HEK293 cells with MLN7243, a small molecule inhibitor of the ubiquitin-activating enzyme UAE1 (Traore et al., 2014), followed by NaAsO_2_ treatment. A 1μM MLN7243 treatment for 2h depletes poly-ubiquitin chains while treatment with either NaAsO_2_ (500 μM for 1h) or the proteasome inhibitor MG132 (50μM, 6h) results in an increase in poly-ubiquitylation (Figure 3B). Despite the greatly reduced level of global poly-ubiquitylation after a 2-hour pre-treatment with 1μM MLN7243, we observed no noticeable impact on SG formation (Figure 3C,D). While this finding strongly suggests that acute protein ubiquitylation is not required for SG formation, it leaves open the possibility that existing ubiquitin chains may have a function in SG assembly. Of particular interest in this regard are lysine 63 (K63)- linked poly-ubiquitin chains, which have been shown to promote the formation of insoluble protein inclusions (Tan et al., 2008) and fibrillar aggregates (Morimoto et al., 2015), possibly by serving as a scaffold for regulated protein aggregation (Pourcelot et al., 2016). To test this hypothesis, we used CRISPR/Cas9-mediated genome editing to generate HEK293 knockout cell lines for UBE2N, which is the E2 enzyme responsible for the majority of K63-linked polyubiquitylation (Hofmann and Pickart, 1999, 2001). We successfully isolated clones in which UBE2N protein levels are reduced to almost undetectable levels by western blot (Figure 3E). Notably, these cell lines are viable with only slightly reduced growth rates, suggesting that K63-linked ubiquitylation is not essential for proliferation of unstressed cells. More importantly, however, lack of UBE2N and the resulting decrease in K63 poly-ubiquitin chains (Figure 3E) have no measurable effect on SG formation (Figure 3F,G). Taken together, these results demonstrate that ubiquitylation is not required for SG formation.

**Figure 3.**
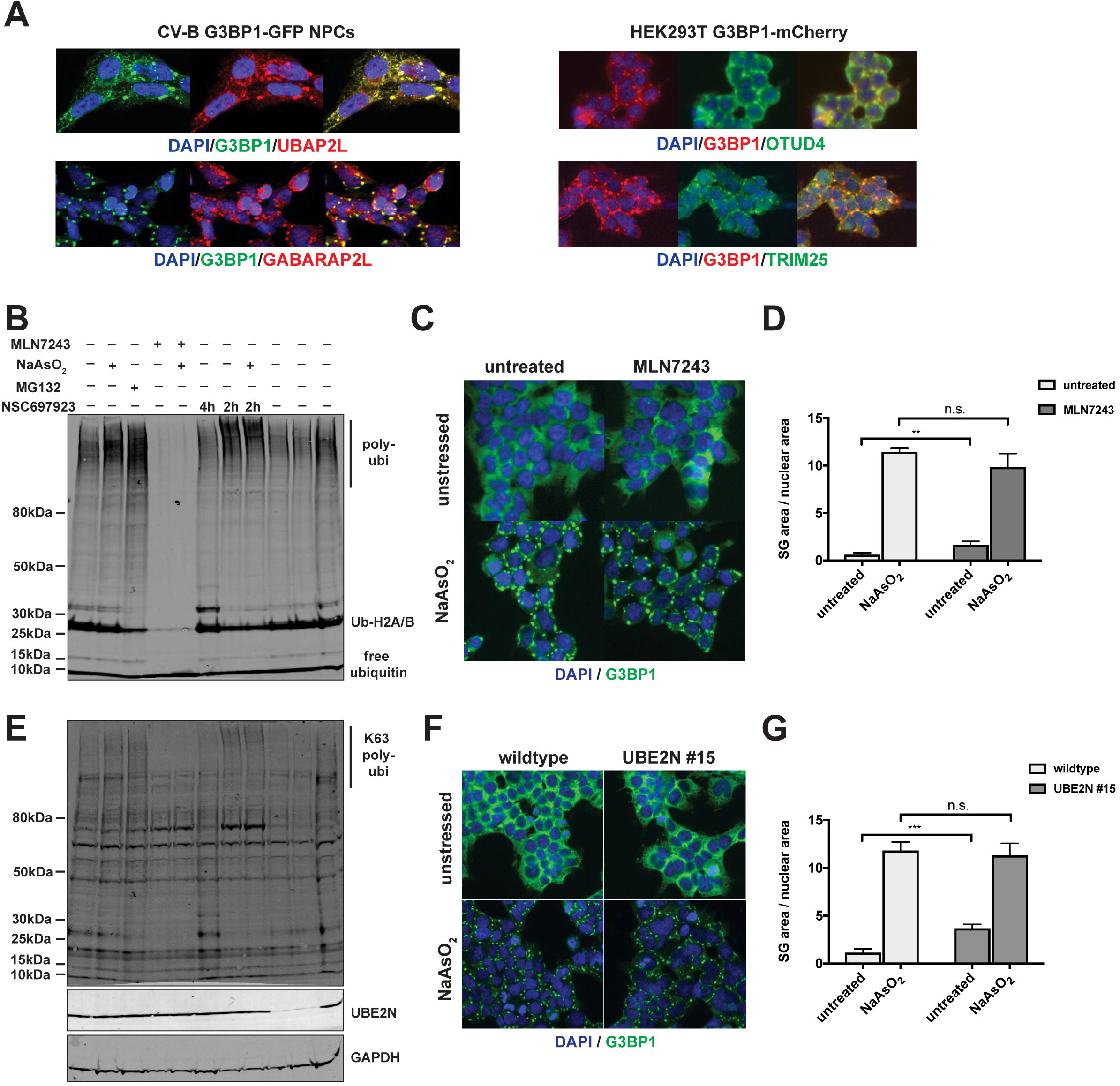
Neither acute ubiquitylation nor non-degradative K63 poly-ubiquitin chains are required for SG formation. **A)** Immunostainings of candidate SG proteins. **B)** Western blot analysis of 293XT cells treated with NaAsO_2_, the proteasome inhibitor MG-132 and the global ubiquitylation inhibitor MLN-7243. Ubiquitin was detected with an antibody against all forms of ubiquitin. **C)** Immunostaining of 293XT cells treated with NaAsO_2_ and the global ubiquitylation inhibitor MLN-7243. **D)** Quantification of SG area in cells treated MLN-7243 and stressed with NaAsO_2_. **E)** Western blot analysis of UBE2N expression and its effect on lysine 63-linked and global poly-ubiquitin chains. **F)** Immunostaining of 293XT cells with and without CRISPR-induced mutations in UBE2N that were treated with NaAsO_2_. **G)** Quantification of SG area in wild type and UBE2N KO cells stressed with NaAsO_2_.

### High-throughput imaging of RBPs reveals cell type- and stress type-specific SG composition

To extend our characterization of SG composition, we focused on the fact that the set of 273 G3BP1-interacting SG proteins identified by our APEX approach is highly enriched for RBPs, with 52% identified as RBPs (Beckmann et al., 2015; Brannan et al., 2016; Gerstberger et al., 2014) compared to 33% (1167/3586) among the background set of all proteins detected in our MS experiments. Given this overrepresentation of RBPs in the SG proteome, we decided to take advantage of a resource of highly validated antibodies directed against >300 human RBPs (Sundararaman et al., 2016) to further characterize the repertoire of RBPs that localize to SGs. For this, we optimized a high-content screening pipeline involving systematic immunofluorescence (IF) labeling of untreated and NaAsO_2_-treated cells (Figure 4A). We first assessed optimal experimental conditions to induce SG formation in different cell types and under distinct stress conditions. To identify RBPs whose localization to SGs is cell type-specific, we conducted parallel screens in three different cell types: HepG2, HeLa and NPCs. Cells were seeded into micro-titer plates and either left untreated or subjected to NaAsO_2_ treatment followed by fixation and IF staining with individual test RBP antibodies and TIA1 as a marker for SGs (Figure 4B). Subsequent high content microscopy and image analysis identified a total set of 77 RBPs that localize to NaAsO_2_-induced SGs induced in at least one of the three tested cell types **(Figure 4C, Table S5**). Over half of these RBPs (42/77) were localized to SGs in all three cell types, with the remaining proteins exhibiting some cell-type specificity (Figure 4B,C). For example, UBAP2L co-localized with SGs in all cell types, while SRSF9, EIF3A and SRP68 were selectively targeted to SGs in HepG2, HeLa or NPCs, respectively (Figure 4D). These results extend our SG compendium and suggest that SGs contain both constitutive RBPs and proteins that exhibit cell-type specific targeting under identical stress conditions.

**Figure 4.**
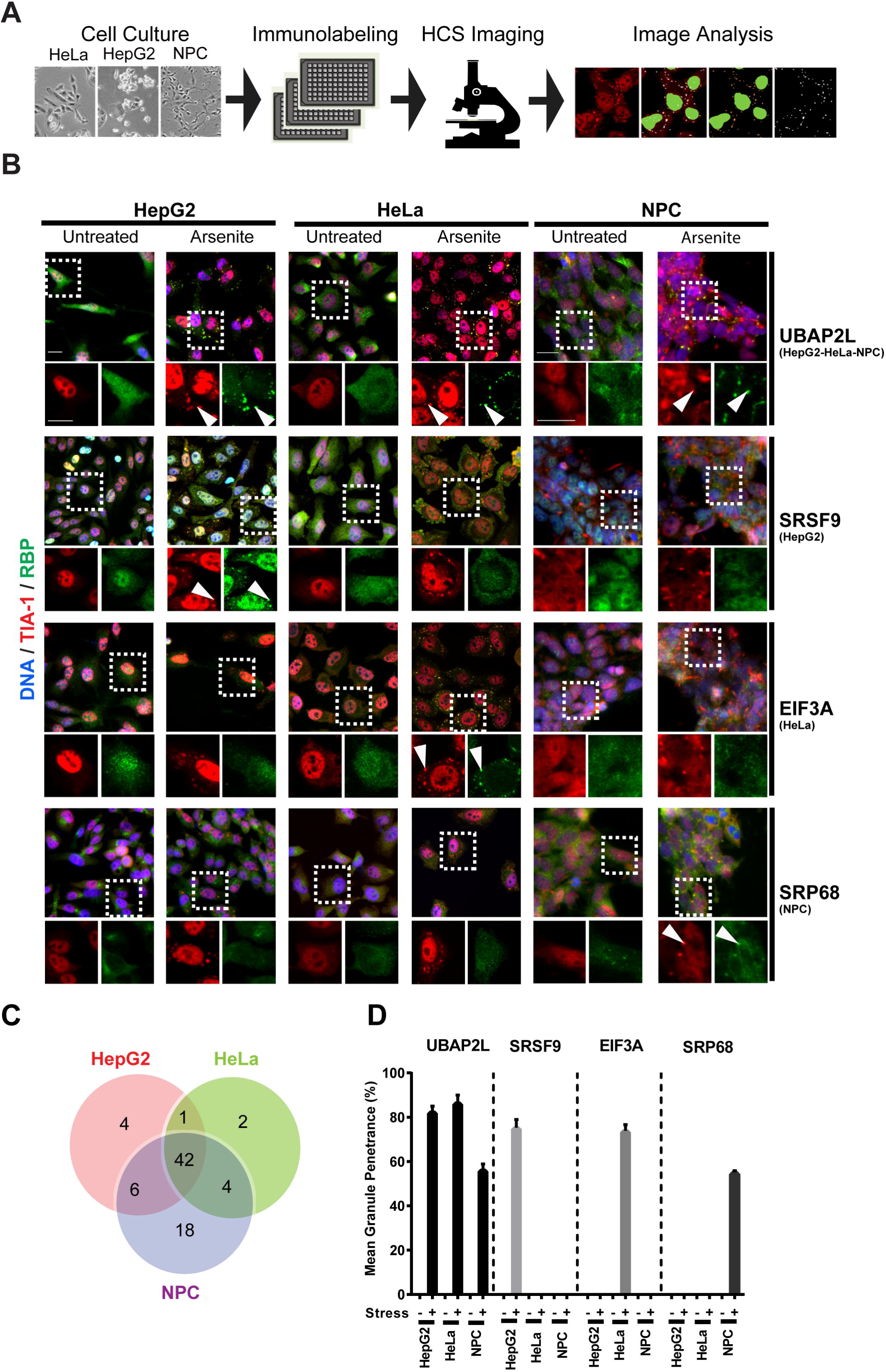
An RBP-centered high-throughput imaging screen identifies cell type-specific SG components. **A) Imaging screen outline to identify RBPs localized to SGs.** HepG2, HeLa and NPC cells were plated within 96 well plates, exposed to specific stresses, and processed for systematic immunofluorescence (IF) using a collection of validated antibodies to human RBPs in combination with the TIA-1 SG marker. High-content imaging was performed to capture images, followed by image analysis and feature quantification using MetaXpress software. The panel on the right shows an example of images obtained with HCS in normal (top panel) versus stressed (bottom panel) cells. **B) Cell-type specificity in RBP localization to SGs.** Examples of RBP localization in untreated and NaAsO_2_-treated HeLa, HepG2 or NPCs. UBAP2L is a common SG hit among the three cell types, SRSF9 is HepG2 specific, EIF3A is HeLa specific, and SRP68 is NPC specific. White arrows indicate examples of RBPs co-localized with the SG marker. In each panel, the indicated insets at the bottom are zoomed views of the same field showing TIA1 (red) and the corresponding RBP (green). Scale bar, 20um. **C)** Venn diagram showing results of imaging screens using 300 validated RBP antibodies in HepG2, HeLa and NPC cell lines. **D)** Mean granule penetrance of proteins with either cell type-independent or cell type-specific SG localization.

Previous studies have shown that several proteins can exhibit stress-dependent variability in SG targeting (Aulas et al., 2017; Kedersha et al., 1999; Tourriere et al., 2003), but this has not been evaluated comprehensively for a large panel of SG-localized proteins. To determine the degree to which SG composition differs depending on the type of stressor, we next performed a parallel screen with our RBP antibody collection in HeLa cells exposed to heat shock (30 min at 42°C), using the same systematic imaging approach (Figure 5A). Similar to our earlier analysis of APEX hits in NaAsO_2_-versus thapsigargin-treated NPCs, we found that the majority of RBPs (40/52) localized to SGs in both stress conditions, while also identifying several proteins that exhibit stress-specificity in SG targeting (Figure 5A-C, **Table S6**). For example, UBAP2L again robustly localized to SGs in both stress conditions, while NOLC1 and SF1 showed stress type-selective SG association in NaAsO_2_-treated or heat-shocked cells, respectively (Figure 5A,C). From this systematic RBP survey, we conclude that approximately one fifth of SG components are modulated differentially depending on the type of stress.

**Figure 5.**
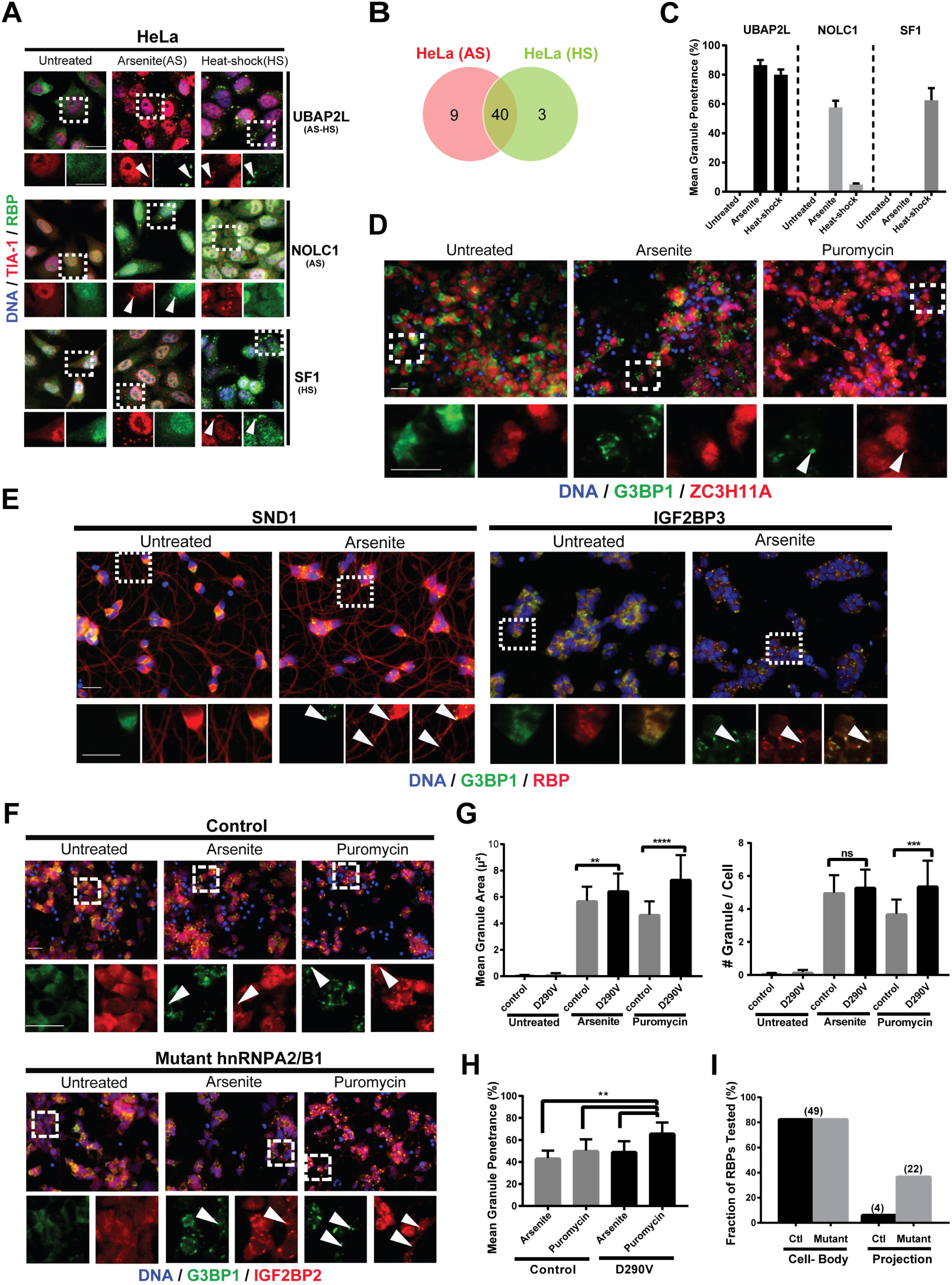
SG composition and subcellular distribution is stress type-dependent. IF images showing examples of RBP localization in untreated, NaAsO_2_ (AS) and heat-shock (HS) treated HeLa cells. UBAP2L is a common hit in both stress condition, NOLC1 is NaAsO_2_-specific and SF1 is heat-shock specific. White arrows indicate co-localization of TIA1 marker with the RBP in SGs in merged views and yellow arrows show TIA1 or RBP, respectively. Scale bar, 20um. **B)** Venn diagram indicating a comparative study of NaAsO_2_ versus heat-shock treated HeLa cells using the same set of antibodies as in Fig 4C. **C)** Mean granule penetrance of proteins with either stress type-independent (UBAP2L) or stress type-specific (NOLC1, SF1) SG localization. **D)** IF images of control iPSC-derived motor neurons treated with either NaAsO_2_ or puromycin to induce SGs. ZC3H11A is co-localized with G3BP1 in puromycin treated cells, but not in NaAsO_2_ treated cells. Upper panels are merged views with lower resolution. In each panel, the indicated insets at the bottom are zoomed views of the same field showing G3BP1 (green) and ZC3H11A (red). **E) Different localization pattern of RBPs in iPSCderived motor neurons (MNs).** IF images of SND1 and IGF2BP3 localization in MN cells (red), which both show cell body staining. SND1 is enriched in neuronal projections and cell body versus IGF2BP3 showing cell body localization. Both RBPs form SGs (arrowheads) in response to stress Upper panels are merged views with lower resolution. In each panel, the indicated insets at the bottom are zoomed views of the same field showing G3BP1 (green) and corresponding RBP (red). **F) IGF2BP2 is co-localized with G3BP1 in neuronal projections in puromycin-treated mutant MN cells.** Control and mutant cells were untreated or stressed with NaAsO_2_ or puromycin, then co-labeled with G3BP1 (green), IGF2BP2 (red) antibodies. Upper panels are merged views with lower resolution. In each panel, the indicated insets at the bottom are zoomed views of the same field showing G3BP1 (green) and IGF2BP2 (red). **G)** Quantification of SG number, area and penetrance in control and hnRNPA2B1 mutant iPS-MNs. **H)** Comparison of RBPs that localized to dendritic SGs in control and hnRNPA2B1 mutant cells. See also Figure S3.

### Stress-induced granules vary in composition and subcellular localization in IPSC-derived motor neurons

The components of SGs as well as the molecular interactions that determine SG dynamics are increasingly implicated in human neurological disorders including ALS (Zarei et al., 2015). As motor neurons (MNs) are the most severely affected cell type in ALS, we characterized the SG-targeting behavior of RBPs in MNs generated by directed differentiation of induced pluripotent stem cells (aspics) (Martinez et al., 2016).This differentiation routinely yields cultures with >90% cells that stain positive for the motor neuron markers Isl-1 and SMI-31 (**Figure S3A**). We first carried out IF staining for 63 (of 77) SG-RBP hits in control iPS-MNs that were either untreated or subjected to stress either by treatment with 250μM NaAsO2 for 60 minutes or with translation inhibitor puromycin, which robustly induces SGs in MNs after a 24h treatment without overt toxicity (Martinez et al., 2016). This analysis identified a collection of 51 RBPs that exhibit co-localization with G3BP1-labelled SGs in MNs treated with either NaAsO_2_ or puromycin (Figure 5D, **Table S7**). Most of these RBPs (49/51) were SG-localized under both conditions, while a few (e.g. ZC3H11A) were selectively targeted in response to puromycin (Figure 5D). Interestingly, in unstressed MNs, we found that several of the tested RBPs exhibited differential localization within cell bodies and dendritic projections (Figure 5E), with 36 of 63 RBPs showing restricted cell body-localization, while 27 of 63 proteins were also detected in neuritis (**Table S7)**. For example, SND1 localized both to cell bodies and projections, while IGF2BP3 was restricted to cell bodies (Figure 5E). In control MNs treated with NaAsO_2_, G3BP1-labeled SGs also demonstrated subcellular specificity, being restricted to cell bodies and mostly devoid from neurites. Both SND1 and IGF2BP3 co-localized with G3BP1-labelled SGs in cell bodies, while SND1 was also present in stress-induced G3BP1-negative granules in dendrites (Figure 5E). These results suggest that stress-induced granules of varying composition form in a subcellular compartment-specific manner in motor neurons.

### SG composition and subcellular distribution are affected in iPSC models of neurodegenerative disease

The inherited forms of neurological disorders such as ALS are often caused by mutations in RBPs known to localize to SGs, such as TDP-43, FUS/TLS and hnRNPA2/B1 (Kim et al., 2013; Kwiatkowski et al., 2009; Sreedharan et al., 2008). To extend our analysis of SG composition and subcellular localization to iPSC-MNs derived from an ALS patient, we performed comparative IF staining on iPSC-derived MNs carrying the ALS-associated D290V mutation in the RBP hnRNPA2B1 (Martinez et al., 2016) (Figure 5F). Image analysis of mutant MNs revealed an increase in both the number and size of G3BP1-postitive SGs in hnRNPA2B1 mutant iPS-MNs compared to control cells (Figure 5G), consistent with Martinez et al. Surprisingly, in addition to the increased propensity to form SGs, we also observed mutation-specific differences in subcellular localization of SGs. In wild type cells, most RBPs localize primarily to SGs in the soma upon puromycin treatment, while in puromycin-treated mutant cells approximately a third of RBPs also localized to SGs in neurites, (Figure 5H) as shown for IGF2BP2 (Figure 5F), as well as IGF2BP1, IGF2BP3, PCBP2, NKRF and FAM120A (**Figure S3B).** These unexpected findings add to the growing body of evidence implicating ALS-causing mutations in RBPs in the abnormal formation and subcellular localization of SGs.

### Cross-comparison with related datasets identifies disease-relevant SG proteins

In combination, our APEX and IF screening approaches identified 314 SG proteins, adding more than 200 proteins that had not previously been associated with SGs. Consistent with what has been reported for known SG proteins, our hits are enriched for RBPs (226/313 = 72.2%, Figure 6A) with a range of RNA-binding domains (Figure 6B) and GO terms associated with RNA metabolism and translational control (Figure 6C). They also contain a significantly higher proportion of amino acid residues in IDRs and LCDs (Figure 6D) than the background proteome, consistent with the hypothesis that liquid-liquid phase separation (LLPS) is one of the drivers of SG assembly. While these observations underscore the specificity of our approach, they do not contribute much to deepen our understanding of how SGs and neurodegeneration are connected. To achieve this, we instead integrated our comprehensive SG proteome with several proteomics datasets relevant to the phenomenon of protein aggregation in neurodegeneration. These include a list of proteins with domains similar to yeast Prion-like domains (PrLD) (March et al., 2016), a set of proteins precipitated by biotinylated isoxazole (Kato et al., 2012), the precipitated SG core proteome (Jain et al., 2016), proteins found to interact with dipeptide repeats derived from the ALS-linked C9ORF72 repeat expansion (Lee et al., 2016; Lin et al., 2016) and proteins found to interact with the ALS-relevant RBPs ATXN2, FUS and TDP-43 (Blokhuis et al., 2016; Freibaum et al., 2010). Lastly, we calculated the proportion of amino acids in LCDs and IDRs for all 1,342 proteins that were detected in at least one of the 14 datasets (**Table S8).** For further analysis, we ranked the proteins in the list by how frequently they occur across all datasets. Notably, only slightly more than one third of proteins (36%, 484/1,342) were detected more than once, with most proteins (64%, 858/1,342) present only in one of the 14 datasets (**Figure S4A**). While these are diverse and distinct interactome datasets that are not expected to overlap completely, the cross-comparison between them nevertheless can provide useful contextual information to the results of each individual study.

**Figure 6.**
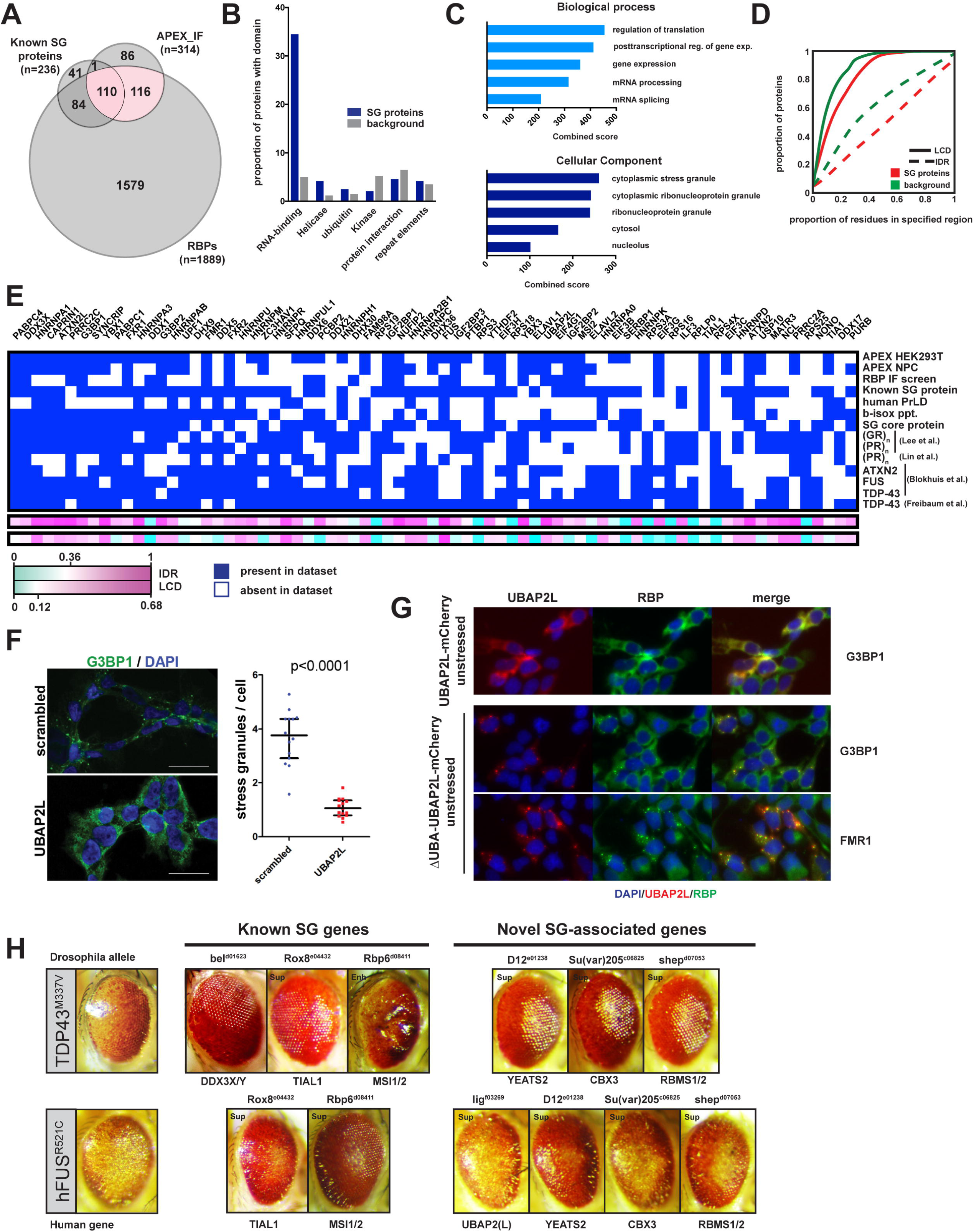
Integrative data analysis highlights potential disease-relevant proteins. **A)** Venn diagram showing overlap between proteins identified in our combined APEX-IF approach, known SG proteins and proteins found to bind RNA. **B)** Distribution of protein domains across SG proteins compared to background. **C)** Gene ontology analysis for 314 APEX-IF hits confirms known functions of SG proteins. **D)** Comparison of the proportion of amino acids in LCDs and IDRs between the 314 SG-APEX hits and background. **E)** Heatmap for the 75 proteins most broadly represented across selected SG and neurodegeneration-relevant datasets. Heatmap indicates whether a protein is present (blue box) or absent (white box) from each dataset and proteins are ranked by the number of datasets they are part of in descending order from left to right. **F)** IF images of G3BP1 staining and quantification of SG number/cell in HeLa cells treated with control siRNA or siRNA targeting UBAP2L. **G)** IF images of 293FITR cells with inducible expression of either a full-length UBAP2L-mCherry fusion protein (top panel) or a truncated UBAP2L-mCherry fusion protein missing the N-terminal UBA domain (middle and bottom panels). Cells were induced with doxyxycline for 24h before fixation. **H)** Images of Drosophila eye degeneration models crossed with the indicated strains. See also Figure S4.

We next focused on the most broadly represented proteins by examining the top 75 proteins in the list in greater detail (Figure 6E). Notably, all of these, and indeed 97% (194/200) of the top 200 proteins, have been identified as RBPs, and most of them contain significantly higher proportions of LCDs and IDRs (Figure 6E) than the background proteome. The list contains many of the best-studied neurodegeneration-linked proteins (e.g. FUS, ATXN2, FMR1, FXR1/2 etc.) and provides a picture that is highly consistent with the emerging view that RBPs in particular are prone to abnormal aggregation in neurodegenerative disease, likely driven or accelerated by LLPS.

Integrating many datasets into a high-level overview also enables prioritizing further candidate genes for follow-up studies in the context of particular diseases or mutations. For example, it was recently shown that cancer-associated mutations in DDX3X, the most highly represented gene across all datasets, likely act by driving increased SG formation (Valentin-Vega et al., 2016). In the context of neurological diseases, DDX3X has so far only been associated with X-linked mental retardation (Snijders Blok et al., 2015). Nevertheless, its strong localization to SGs, broad representation across neurodegeneration-relevant datasets and the fact that it is already a target of interest for the development of small molecule inhibitors (Bol et al., 2015) make DDX3X an interesting candidate for follow-up studies. Another example is UBAP2L, which had not previously been associated with either neurological diseases or stress granules. We found UBAP2L to be consistently among the most highly enriched proteins by SG-APEX and it robustly localized to SGs across all cell types and stress conditions tested (Figures 4B, 5A). Strikingly, our depletion of UBAP2L by siRNA in HeLa cells almost completely abolished NaAsO_2_-induced SG formation (Figure 6F). Surprisingly, the N-terminal ubiquitin-associated (UBA) domain of UBAP2L was not required for its localization to SGs, in contrast to the ubiquitin-binding domain of HDAC6 (Kwon et al., 2007). Instead, inducible expression of a UBAP2LmCherry fusion protein lacking the UBA domain ( ΔUBA_UBAP2L-mCherry) led to widespread formation of aggregates that also stained positive for the SG-associated proteins G3BP1, FMR1 and ELAVL1 (Figure 6G, **Figure S4B**). In contrast, full-length UBAP2L-mCherry only showed a slight tendency to form spontaneous aggregates in the absence of stress (Figure 6G). Although UBAP2L localizes to SGs independently of its ubiquitin-associated functions, it is possible that the UBA domain could regulate its tendency to aggregate through interactions with ubiquitylated substrates. Our findings establish UBAP2L as a highly disordered (**Figure S4C**) and essential regulator of SG formation and a candidate for further studies into mechanisms of protein aggregation and its possible role in neurodegenerative disease.

### Known and novel SG proteins can modify FUS and TDP-43 toxicity in a fly model of neurodegeneration

To further evaluate if components of SGs provide insights into evolutionarily conserved links to neurological disease, we compared selected SG proteins to a resource of *Drosophila* genes shown to modify eye degeneration phenotypes caused by overexpression of mutant versions of either FUS or TDP-43 (unpublished data, Lanson et al., 2011; Periz et al., 2015; Ritson et al., 2010). Indeed we found that several known (DDX3X/Y, TIAL1 and MSI1/2) and previously unknown (YEATS2, CBX3, RBMS1/2 and LIG/UBAP2) SG proteins were able to suppress TDP-43- and FUS-mediated toxicity (Figure 6H). These examples strongly demonstrate that our newly established SG-proteome resource is valuable in identifying candidate genes with putative roles in modulating or precipitating neurological dysfunction.

### Protein-protein interaction analysis reveals a dense steady-state interaction network of SG proteins

Lastly, to provide a global view of protein-protein interactions between SG proteins and neurodegeneration-linked proteins, we generated protein-protein interaction (PPI) networks for the 426 SG-associated proteins described above. We retrieved only experimentally validated direct protein-protein interactions from the mentha (Calderone et al., 2013), MINT (Licata et al., 2012) and IntAct (Orchard et al., 2014) databases through the Proteomics Standard Initiative Common QUery InterfaCe (PSICQUIC) web portal (del-Toro et al., 2013) and merged them with the BioPlex protein interaction dataset (Huttlin et al., 2017; Huttlin et al., 2015). The resulting global PIN contains 14,352 nodes and 102,551 non-redundant edges between nodes and the SG-PIN subnetwork with 426 nodes and 1129 edges was selected and visualized using Cytoscape (Shannon et al., 2003) (Figure 7A).

**Figure 7.**
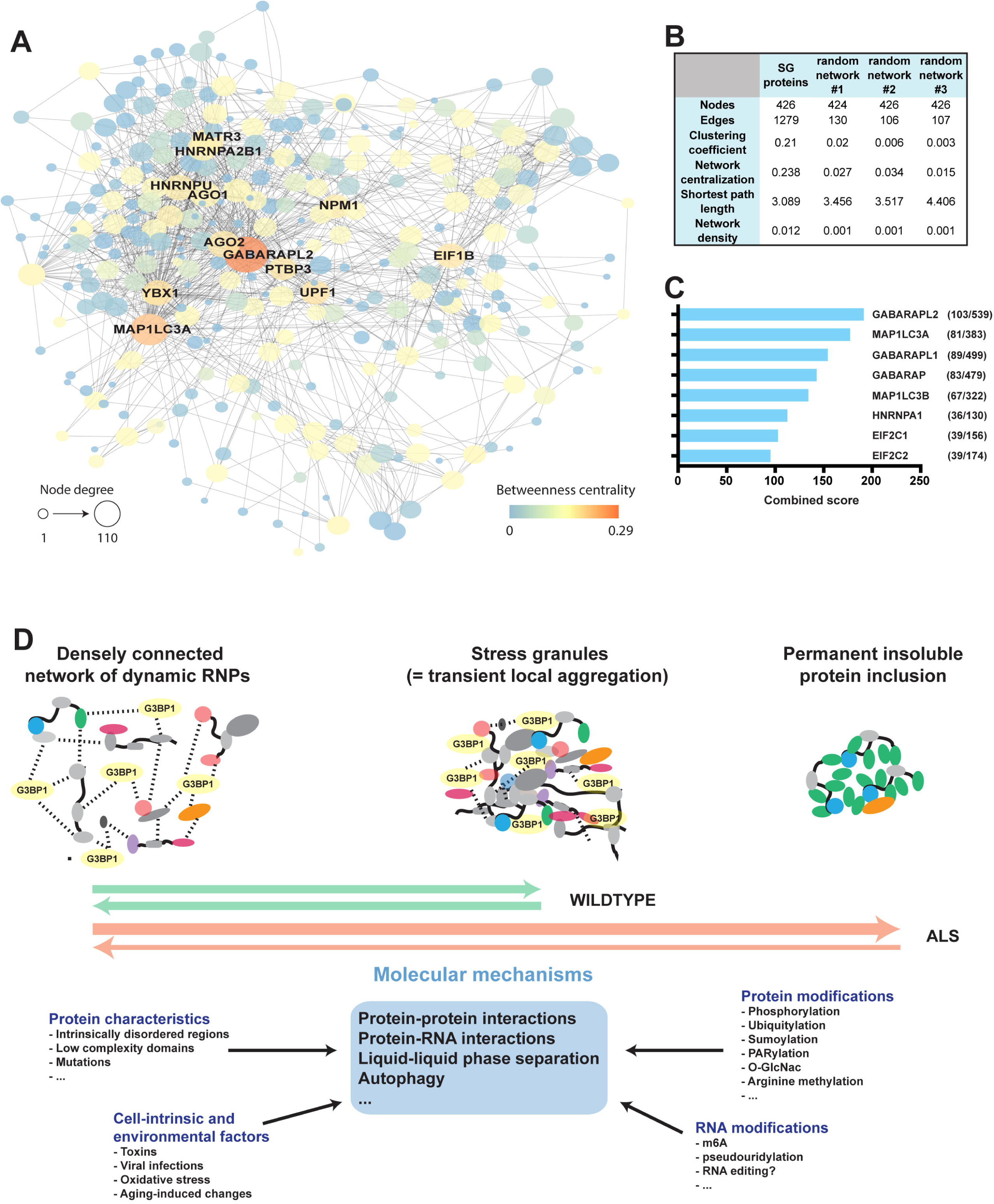
SG proteins form a pre-existing protein interaction network that enables rapid formation and resolution of SGs. **A)** Protein interaction network (PIN) of 426 proteins identified as APEX-IF hits or previously shown to associate with SGs. Network was visualized in Cytoscape using a force-directed layout. **B)** Common network parameters for the SG-PIN compared to three PINs from a randomly selected equal number of nodes. **C)** Ranked list of proteins with the greatest connectivity to the SG-PIN as determined by the Enrichr gene enrichment analysis tool. **D)** Model of the relationship between normally functioning, dynamic RNPs, transiently aggregating SGs and permanently aggregated pathological protein inclusions.

One of the most interesting findings from our APEX data was the observation that the G3BP1 interactome is remarkably stable upon exposure to stress, suggesting that the rapid formation of SGs is primed by pre-existing protein interactions. In strong support of this conclusion, the SG-PIN built from a combination of multiple independent and unbiased PPI datasets acquired in the absence of cellular stress shows that SG proteins indeed form a very dense interaction network. By comparison, networks built from an equal number of randomly selected nodes have on average about 90% fewer edges and score significantly lower on other common parameters used to describe network topology and connectivity such as clustering coefficient, network centralization and shortest path length (Figure 7B).

### The SG protein interaction network highlights autophagy as a central mechanism in SG biology

Intriguingly, the two most central nodes of the SG-PIN are GABARAPL2 and MAP1LC3A, which are both small ubiquitin-like modifiers of the ATG8 protein family directly implicated in different steps of autophagy (Ichimura et al., 2000) (Figure 7A). In addition, an analysis of protein-protein interaction (PPI) hubs through the Enrichr analysis tool (Chen et al., 2013; Kuleshov et al., 2016) reveals that the 5 proteins with the highest connectivity to the SG-PIN are all ATG8 family members (Figure 7C). To underscore the significant enrichment of these interactions, while only about 3% (426/14352) of all proteins in the dataset are SG-associated, 19.1% (103/539) of all GABARAPL2 interactions and 21.1% (81/383) of all MAP1LC3A interactions are with SG proteins (Figure 7C). The PPIs of autophagy-related proteins have previously been studied extensively (Behrends et al., 2010) and shown to overlap with disease-associated PINs (Wang et al., 2017). Our results clearly support the previously described function for autophagic processes in SG biology (Buchan et al., 2013). Indeed, they further indicate that pre-formed interactions with autophagy proteins may allow for efficient clearance of SG-associated proteins even in the absence of cellular stress.

## Discussion

### APEX2-mediated proximity labeling allows for the proteomic characterization of RNP granules

In this study we expand the application of APEX proximity labeling to highly dynamic, non-membranous RNP granules that are not restricted to any particular subcellular localization. By using a CRISPR/Cas9-engineered cell line in which the core SG component G3BP1 has been endogenously tagged with an APEX2-GFP fusion protein, we were able to identify dozens of proteins not previously known to associate with SGs. Some of these are family members of previously identified SG-associated proteins (e.g. YTHDF3) or are known to interact directly with a variety of known SG proteins (e.g. CSDE1). However, many of the proteins had not previously been connected to SGs. Through cross-comparison of our SG-APEX dataset with other SG datasets (Jain et al., 2016) as well as PPI datasets relevant to neurodegeneration (Blokhuis et al., 2016; Freibaum et al., 2010; Lee et al., 2016; Lin et al., 2016) we confirm that the APEX approach as the only *in vivo* labeling method is both sensitive and specific in identifying SG proteins. Nevertheless, it may be possible to further address the specific challenges of RNP granules, such as their potentially tightly structured molecular architecture by combining and integrating proximity labeling proteomics data from several different APEX2-tagged RNP granule proteins. Not only should this reduce the rate of false positive and negative hits but careful selection of APEX2 fusion partners should also enable a higher resolution characterization of subtypes of closely associated RNP granules, such as P-bodies and SGs.

### SG composition shows higher degree of cell-type and stress-specificity than previously described

In addition to HEK293T cells, we also performed G3BP-APEX proximity labeling in CRISPR/Cas9-engineered iPSC-derived neural progenitor cells, underscoring that this method is applicable to a wide range of relevant cell types. While cell-type specific differences in SG composition had been previously described for individual proteins and cell types, we provide what is to our knowledge the first systematic characterization of how SG composition varies across cell types and stress conditions. We find that while the majority of SG proteins are localize robustly across conditions, there are clear differences, in particular between different cell types. Differential protein expression levels across cell types likely contribute, but cannot by themselves explain all the variation we observed. It will be interesting to further investigate the basis of the cell type specificity by examining the presence of different protein isoforms, as well as cell-type specific protein-protein and protein-RNA interaction networks. These differences may underlie cell-type vulnerability in neurological disease.

### A model of SGs as part of a regulated equilibrium of RNP interactions that is disrupted in neurodegeneration

By employing different experimental designs to probe the G3BP1 interactome in both stressed and unstressed cells, we showed that many interactions between SG proteins already exist in unstressed cells. This observation was further supported by our PPI analysis of SG proteins, which showed that SG proteins form a dense network of PPIs in unstressed cells. Remarkably, it appears that SG proteins even at steady state are tightly surveilled by the cellular autophagy system, suggesting that an efficient mechanism for disassembly and subsequent clearance of SGs is programmed into the system. It is moreover striking that the overwhelming majority of proteins that are prominently found in SGs and are broadly represented in neurodegeneration-associated datasets have the capacity to bind RNA, and closely interact with each other, either directly or likely through common RNA targets. This suggests that for SGs in particular, RNA is likely to play a more important role than just acting as cargo that is to be sequestered from translation. Indeed, free mRNA has been shown previously to act as a scaffold for SG formation (Bounedjah et al., 2014), presumably by allowing its bound RBPs to multimerize. The resulting close proximity of large amounts of SG-RBPs allows these proteins, many of which contain high proportions of amino acids in LCDs or IDRs, to undergo LLPS that further drives SG growth. However, it will be interesting to investigate whether mRNA molecules may have a more active role in SG formation, for example by regulated RNA-RNA interactions or by active participation of RNA in LLPS. The observation that the binding profile across transcript regions of the critical SG proteins G3BP1 and TIA1 shifts significantly upon stress (unpublished data) suggests that RNA may play more than just a passive role in SG formation.

Based on our findings, we present a refined model for SG dynamics that views SGs as a highly evolved system by which cells can rapidly respond to stress conditions, while at the same time ensuring that mechanisms are in place to quickly and efficiently resume normal function upon return to more favorable conditions. (Figure 7D). Most importantly, we view SGs not as a distinct cellular organelle that forms completely *de novo* upon exposure to stress, but rather as a transient and tightly controlled extreme in a highly dynamic landscape of protein-protein and protein-RNA interactions. Much attention has recently been focused on the role of LLPS in both transient and permanent protein aggregation, especially since many ALS-causing mutations in SG-associated RBPs are thought to increase the propensity of these proteins to aggregate. This is likely to be an important factor in the aberrant SG formation and distribution we observe in hnRNPA2B1 mutant ALS patient motor neurons.

However, we are only just beginning to understand that the normal PPI landscape (both RNA-dependent and RNA-independent) is dependent on a complex system of regulatory post-translational protein modifications. Examples include the regulation of G3BP1 self-aggregation by phosphorylation (Kedersha et al., 2016), modification of SG proteins by autophagy mediators to allow clearance of aggregates (Buchan et al., 2013) and the complex multi-layered regulation of SUMO-controlled protein interaction through acetylation of the SUMO regulator itself (Ullmann et al., 2012). Moreover, arginine methylation in LCDs of some SG proteins such as FUS/TLS has been shown to modulate their subcellular distribution and toxicity (Tradewell et al., 2012). Lastly, there are several less well-understood SG-associated PTMs such as PARylation (Leung et al., 2011), modification by O-GlcNAc (Ohn et al., 2008) and age-related changes in regulation of oxidative stress and protein turnover.

Importantly, a model in which sustained disruption of one or more of these mechanisms regulating the equilibrium of RNPs can ultimately lead to the formation of insoluble protein aggregates could explain how a variety of seemingly unrelated mutations in multifunctional RBPs converge to cause neuronal cell death through a common pathway. It is in this context that we can leverage data from genetic interaction screens such as the fly modifiers we describe. By understanding the normal function of the genes identified as modifiers of neurotoxicity, we can further elucidate the mechanisms that are most critical in connecting SGs and disease.

As a result, understanding the complex network of regulatory modifications may hold a vast potential for identifying new strategies for targeted therapies of protein aggregation in a wide range of neurodegenerative diseases. It will be critical however, that these efforts can be finely tuned and are designed to specifically target pathological protein aggregation, such as not to disrupt the dynamic RNP landscape in healthy cells.

## Experimental Procedures

### Cell Culture

HEK293T and HeLa cells were maintained in DMEM and HepG2 cells in Hyclone growth medium both supplemented with 10% fetal bovine serum and 1% penicillin/streptomycin. For SILAC experiments, DMEM without L-arginine and L-lysine (Pierce catalog no. PI88420) was supplemented with 10% dialyzed FBS (Pierce, PI88440), penicillin/streptomycin, and 0.4mM and 0.8mM, respectively, of either unlabeled L-Lysine:HCL and L-Arginine:HCl (Sigma, cat no. L8662 and A6969) or isotopically labeled L-Lysine: 2HCl (^13^C_6_, ^15^N_2_) and L-Arginine:HCl (^13^C_6_, ^15^N_4_) (Cambridge Isotope laboratories, cat no. CNLM-291 and CNLM-539). Both heavy and light medium were additionally supplemented with 200mg/ml L-Proline (Sigma, cat no. P5607).

Neural progenitor cells were derived from iPSCs grown in mTeSR1 (Stem Cell Technologies) as described in Reinhardt et al., 2013 and grown in medium consisting of DMEM/F12+Glutamax, 1:200 N2 supplement, 1:100 B27 supplement, penicillin/streptomycin (Life technologies) and 100μM ascorbic acid (Sigma, A4544), supplemented with 3μM CHIR99021 (CHIR) and 0.5μM Purmorphamine (PMA) (Tocris, cat no. 4423 and 4551). For SILAC experiments, DMEM/F12 without L-arginine and L-lysine (Pierce catalog no. PI88215) was used instead of regular DMEM/F12 and supplemented with 0.7mM and 0.5mM, respectively, of either unlabeled L-Lysine:HCL and L-Arginine:HCl (Sigma, cat no. L8662 and A6969) or isotopically labeled L-Lysine: 2HCl (^13^C_6_, ^15^N_2_) and L-Arginine:HCl (^13^C_6_, ^15^N_4_) (Cambridge Isotope laboratories, cat no. CNLM-291 and CNLM-539). Motor neurons were differentiated from iPSCs as described in Martinez et al., 2016. At day 18 of differentiation, cells were dissociated and either plated directly for continued differentiation or optionally expanded in motor neuron progenitor (MNP) medium as described in (Du et al., 2015). For imaging, cells were dissociated again at day 23 and plated into 96-well plates serially coated with 0.001% (=0.01mg/ml) poly-D-lysine (Sigma, P6407) and poly-L-ornithine (Sigma, P3655) followed by 20ug/ml laminin (Life technologies, 23017015). For UBAP2L overexpression experiments, full-length and ΔUBA-UBAP2L-mCherry fusion constructs were cloned into pLIX_403 (gift from David Root, Addgene plasmid # 41395) and packaged into lentiviral particles. MNPs were transduced and selected with 2μg/ml puromycin (Life technologies, A1113803) for 7 days starting 2 days post-transduction. Expression was induced by adding 100ng/ml doxyxycline for 24 hours. To induce SG formation, cells were treated with 250μM (NPCs, MNs) or 500μM (HEK293T, HeLa, HepG2 cells) sodium arsenite for 30min (HeLa cells) or 1h (NPCs, MNs, HEK293T and HepG2 cells). Alternatively, SG formation was induced by treatment with 10ug/ml puromycin for 24h (MNs), 500nM thapsigargin (NPCs) or by heat shock for 30min at 45°C (HeLa, HepG2 cells).

### APEX-mediated biotinylation

HEK293Ts and NPCs were grown in heavy or light SILAC medium for at least 5 passages prior to APEX labeling and isotope label incorporation efficiency was confirmed to be above 98%. Cells were seeded in 10cm culture dishes one day prior to labeling to be ~80% confluent the following day and either left unstressed or treated with either 250μM (NPCs) or 500μM (HEK293T) arsenite or 500nM thapsigargin for 1h at 37°C. 500μM biotin-phenol (BP) was added to the medium at the same time as stressors except for the no-substrate control samples. APEX labeling was performed by adding hydrogen peroxide to a final concentration of 1mM for 60 seconds before quenching the biotinylation reaction by adding Trolox ((+/-)-6-Hydroxy-2,5,7,8-tetramethylchromane-2-carboxylic acid, Sigma 238813) and sodium L-ascorbate (Sigma A4034) to a final concentration of 5 and 10mM, respectively. Samples were washed once with cold PBS, collected using cell scrapers, pelleted for 3min at 300g and immediately suspended in cold lysis buffer (8M urea, 150mM NaCl, 20mM Tris-HCl pH 8.0, Protease Inhibitor Cocktail Set III, EDTA-Free (EMD Millipore, cat no. 539134), 5mM Trolox and 10mM sodium Lascorbate). Samples were sonicated and cleared by centrifugation at 12000rpm for 10min at 4°C. Protein concentration was determined using by 660nm protein assay (Pierce, PI22660) and equal amounts of protein from corresponding light and heavy labeled samples were mixed for a total of 2-4mg of protein. Samples were diluted to 2M urea by adding 3 volumes of 150mM NaCl, 20mM TrisHCl pH 8.0 with protease inhibitors and quenchers. For affinity purification, ~100ul of streptavidin magnetic beads (Pierce, PI88817) were washed once in 2M urea buffer, resuspended directly in the sample, incubated for 2h at room temperature and washed 8 times in 2M urea buffer. Samples were digested directly off the beads and processed as described in Gendron et al., 2016).

### Data analysis

Log2H/L ratios for each protein detected after streptavidin IP samples were input-normalized by subtracting the mean log2H/L ratio of that protein in the corresponding triplicate input samples. For those proteins for which no input data was available, normalization was carried out using the median log2H/L ratio of all proteins in the corresponding triplicate input samples. For hit identification, proteins in each dataset were ranked by descending log2H/L ratios and the fraction of known SG proteins in a rolling window (size=200) was calculated. A cutoff was determined to be the point at which the frequency of known SG proteins fell below 1.5 times (HEK293T cells) or 2 times (NPCs) the background frequency.

### Immunofluorescence, imaging and image analysis

Cells were fixed for 20 min in 4% formaldehyde, 1X PBS, followed by permeabilization for 10 min with 0.5% Triton, 1X PBS. Cells were rinsed with 1X PBS and blocked with blocking buffer (1X PBS, 2% BSA, 0.02% Triton). Cells were incubated with the primary antibodies against stress granule marker like TIA1 (TIA1, dilution 1:100, cat.# sc-1751,Santacruz) and antibodies against RBPs [Table S2,(Sundararaman et al., 2016)] diluted in blocking buffer for 2 hour at room temperature or overnight at 4°C. Then, the cells were thoroughly washed with 1X PBS, 0.2% Tween 20, and incubated for 2 hour with secondary antibodies (Alexa Fluor 647, cat. # A21447, Alexa Fluor 594, cat. # A21207, Life technologies and Alexa Fluor 488, cat. # 111-546-144, JacksonImmuno, dilution 1:500) diluted in blocking buffer. Cells were washed, incubated for 5 min with DAPI and washed again. Cells were stored in the dark at 4°C in 1X PBS or 50% glycerol/PBS for long-term storage. All images were taken using high content screen microscopy, ImageXpress Micro. MetaXpress v3.1 software was used for all image analysis and quantifications were carried out using an in-house script.

### *Drosophila* genetics

Flies were reared on standard yeast-agar-cornmeal medium and crosses were performed at 25°C. *Drosophila* transgenic strains carrying GAL4 inducible human ALS disease causing alleles of FUS/TLS and TDP-43 were previously described (Lanson et al., 2011; Ritson et al., 2010). Standard genetic procedures were used to generated the *GMR-GAL4*/*CyO*, *tub-GAL80*; *UAS-FUS-hR521C*/*TM6B, Tb* and *GMR-GAL4*, *UAS-TDP-43-hM337V*/*CyO*, *tub-GAL80* transgenic strains (Periz et al., 2015). *Drosophila* strains containing the Exelixis insertional disruptions are publically available from the Department of Cell Biology, Harvard Medical School include Rox8^e04432^, Rbp6^d08411^, lig^f03269^, CG2889^d07154^, *D12*^e01238^, *Su(var)205*^c06825^ and *shep*^d0705**3**^. The dominant effect of the introduction of these inserts on degenerative eye phenotypes of *GMR-GAL4*; *UAS-FUS-hR521C* and *GMR-GAL4*, *UAS-TDP-43-hM337V* was assessed two weeks after the crosses were performed. Qualitative changes in pigmentation, ommatidial structure and glossiness phenotypes were monitored for enhancement or suppression.

### Protein interaction network analysis

To retrieve protein interaction data and build protein-protein interaction networks, we queried the Proteomics Standard Initiative Common QUery InterfaCe (PSICQUIC) web portal (http://www.ebi.ac.uk/Tools/webservices/psicquic/view/main.xhtml) for PPI data form the mentha, IntAct and MINT databases. We restricted results to only human interactors that had been experimentally validated in AP-MS experiments (i.e. search terms MI:0006: anti bait coimmunoprecipitation and MI:0007: anti tag coimmunoprecipitation). The resulting data were combined with the most recently available dataset based on AP-MS interactions of ~5000 bait proteins from the Bioplex website (http://bioplex.hms.harvard.edu). We used Cytoscape to visualize the resulting PPI dataset consisting of 14,352 nodes and 102,551 non-redundant edges. We extracted PPI data for 426 SG proteins and used the Prefuse Force Directed Layout to visualize the network. The internal Cytoscape Network Analyzer plugin was used to calculate and visualize network parameters.

### Protein domain and gene ontology analysis

Domain analysis was done by retrieving PFAM domains through the NCBI Conserved Domains Database (https://www.ncbi.nlm.nih.gov/Structure/cdd/cdd.shtml). Low complexity domains and intrinsically disordered regions were calculated as previously described (Beckmann et al., 2015; Conrad et al., 2016). Gene ontology enrichment analysis and PPI hub analysis was performed through the Enrichr Gene Ontology enrichment tool (http://amp.pharm.mssm.edu/Enrichr/) (Chen et al., 2013; Kuleshov et al., 2016).

## Author contributions

Conceptualization, S.M. and G.W.Y; Methodology, S.M., E.J.B., E.L. and G.W.Y.; Formal Analysis, S.M., W.J. and E.J.B.; Investigation, S.M., S.S., K.S., R.M., E.L., F.K., M.W.K., A.S., K.G.G and M.D.L.; Writing – Original Draft, S.M.; Writing – Review & Editing, S.M., E.J.B., E.L., and G.W.Y; Visualization, S.M. and S.S.; Supervision, S.A-T., E.J.B., E.L., G.W.Y., Funding Acquisition, E.J.B., E.L. and G.W.Y.

## Acknowledgements

We acknowledge members of the G.W.Y. lab, particularly Tomas Bos, Kristopher Brannan and Isaac Alexander Chaim for critical comments. K. G. Guruharsha, Maria D. Lalioti and Spyros ArtavanisTsakanos also provided valuable comments on the manuscript. S.M is supported by a postdoctoral fellowship from the Larry L. Hillblom Foundation (#2014-A-027-FEL). S.S. was supported by the IRCM Angelo Pizzagalli fellowship. G.W.Y. is an Alfred P. Sloan Research Fellow and E.L. is an FRQS Junior 2 Scholar. Work in the Lécuyer was supported by an ALS Canada/Brain Canada Discovery Grant. This work was partially supported by grants from the ALS Association to G.W.Y.

